# Pre- and post-sequencing recommendations for functional annotation of human fecal metagenomes

**DOI:** 10.1101/760207

**Authors:** Michelle L. Treiber, Diana H. Taft, Ian Korf, David A. Mills, Danielle G. Lemay

**Author notes:** Corresponding author Email addresses: DGL.

## Abstract

**Background:** Shotgun metagenomes are often assembled prior to annotation of genes which biases the functional capacity of a community towards its most abundant members. For an unbiased assessment of community function, short reads need to be mapped directly to a gene or protein database. The ability to detect genes in short read sequences is dependent on pre- and post-sequencing decisions. The objective of the current study was to determine how library size selection, read length and format, protein database, e-value threshold, and sequencing depth impact gene-centric analysis of human fecal microbiomes when using DIAMOND, an alignment tool that is up to 20,000 times faster than BLASTX.

**Results:** Using metagenomes simulated from a database of experimentally verified protein sequences, we find that read length, e-value threshold, and the choice of protein database dramatically impact detection of a known target, with best performance achieved with longer reads, stricter e-value thresholds, and a custom database. Using publicly available metagenomes, we evaluated library size selection, paired end read strategy, and sequencing depth. Longer read lengths were acheivable by merging paired ends when the sequencing library was size-selected to enable overlaps. When paired ends could not be merged, a congruent strategy in which both ends are independently mapped was acceptable. Sequencing depths of 5 million merged reads minimized the error of abundance estimates of specific target genes, including an antimicrobial resistance gene.

**Conclusions:** Shotgun metagenomes of DNA extracted from human fecal samples sequenced using the Illumina platform should be size-selected to enable merging of paired end reads and should be sequenced in the PE150 format with a minimum sequencing depth of 5 million merge-able reads to enable detection of specific target genes. Expecting the merged reads to be 180-250bp in length, the appropriate e-value threshold for DIAMOND would then need to be more strict than the default. Accurate and interpretable results for specific hypotheses will be best obtained using small databases customized for the research question.

## Background

Measurement of the human fecal microbiome can provide a snapshot of the functional capacity of the microbes of the distal gut. Previous studies have used methods in which a particular marker gene, e.g. 16S rRNA gene, is amplified and sequenced to estimate the taxonomy of microbes present [1]. A method, PICRUSt (phylogenetic investigation of communities by reconstruction of unobserved states), has been developed to predict the functional composition of metagenomes from 16S rRNA marker data and reference genomes [2]. However, in a comparison with full metagenomes, PICRUSt still missed a large percentage of genes known to be present and predicted many genes not found in adequately sequenced metagenomes [3]. The current golden standard remains to sequence all of the DNA present in the sample—the metagenome, also known as the “shotgun” metagenome—and to directly map DNA sequences to protein or gene family databases to infer functional capacity.

One common way of functionally annotating a shotgun metagenome is to first assemble the reads into longer fragments of DNA called contigs, then predict open reading frames in the contigs and map these ORFs to gene family databases. This is valuable because it can provide genomic context for individual genes. However, when estimating the functional capacity of a community, metagenomics assembly biases in favor of the assembled contigs which have less fragmented gene sequences. This means that abundance estimates are biased in favor of the functions of the most abundant organisms. Also, identical genes may exist in multiple species, making their origin unresolvable with short reads. Assembly has been shown to be detrimental to obtaining true gene abundance estimates, as in the case of antibiotic resistance genes [4]. If the objective of the study is to assess the overall functional capacity of a community, it may not be necessary to put sequences in the context of their individual genomes.

Different bioinformatics strategies have been suggested to functionally annotate shotgun metagenomes (reviewed in [5–7]). Regardless of the bioinformatics strategy used, a fundamental step is the ability to detect genes-of-interest in short read sequencing data now commonly produced via Illumina platforms. At the time of this writing, common formats are single read 50bp or 100bp (SR50, SR100) and paired end 100bp and 150bp (PE100, PE150). Gene detection in short read sequences is impacted by decisions in both the pre- and post-sequencing phases. Pre-sequencing decisions include choices of size-selection during library preparation, read length and format (single or paired end), and sequencing depth. Post-sequencing decisions include selection of pre-processing steps, a reference database, and a mapping strategy. While some guidance has been provided on sequencing [8], recommendations are based largely on practices that best inform taxonomy, rather than function. Metagenome studies have generally been designed with microbial ecology (e.g. taxonomy) in mind as the foremost goal with functional annotation as an afterthought. As a result, pre-sequencing decisions for the estimation of taxonomy may be weighted towards creating non-overlapping paired ends of reads that maximize the ability to fully assemble the metagenome. However, tools that map reads to protein databases, such as BLASTX [9] and DIAMOND [10], are not able to leverage non-overlapping pairs of reads. A systematic assessment of the annotation of short reads demonstrated the general utility of homology-based mapping with the caution that read length, phylogeny, and database coverage impacts accuracy [11]. Since that important work which utilized BLAST [11], a protein alignment tool called DIAMOND was developed which is 20,000 times faster than BLASTX [10]. We therefore sought to identify appropriate e-value thresholds for functional annotation of shotgun metagenomics using DIAMOND for read lengths commonly available via the popular Illumina platform in the current study.

Another important limitation of prior work is that it has not been clear how pre- and post-sequencing decisions impact the ability to quantify abundances of genes with verified functionality. Databases of genes, gene families, or orthologous groups are often based on predictions or homology with few entries being verified for functionality. For example, the carbohydrate-active enzymes database (CAZy) requires only one member of each gene family to be biochemically characterized with the remaining members assigned based on amino acid sequence similarity [12]. It is possible for enzymes to be assigned to more than one family and for different enzymes in the same family to act on different substrates.

In the current study, we evaluated various pre- and post- sequencing choices. We first assembled a database of protein sequences for which the function has been experimentally verified and then used this database of “knowns” to generate simulated metagenomes consisting of sequence reads of proteins with verified functionality. We then used these simulated metagenomes to investigate how read length, e-value threshold, and the choice of protein database impacted our ability to detect sequences of known abundance. Next, we used real human stool metagenomes from two different studies to investigate strategies for paired end reads when most reads don’t overlap. Finally, we sub-sampled deeply sequenced fecal metagenomes to determine the minimum sequencing depth needed to quantify the abundance of the genes of specific enzymes.

## Results

### Evaluation of the effect of read length

We first constructed a database containing only protein sequences with experimentally verified functions (see Methods). Metagenomes were then simulated using sequences from this database with increasing proportions of a target enzyme, beta-galactosidase, with the non-beta-galactosidase reads being derived from other sequences in our known protein database. One hundred metagenomes were simulated with five different read lengths of 50 bp, 100 bp, 150 bp, 200 bp, and 250 bp for 500 total metagenomes. For each read length, the 100 simulated metagenomes were mapped to a custom beta-galactosidase database, the NCBI RefSeq database [13] or the SEED database [14] using DIAMOND with the “sensitive” flag and default e-value threshold. With a read length of 50 bp, only the custom beta galactosidase database, from which the beta-gal metagenomic reads were simulated, performed well with both high true positive rates and low false positive rates (**Figure 1**). True positive rates were expressed as the proportion of reads known to originate from the target that were correctly identified as the target. False positive rates would ordinarily be expressed as the proportion of reads not originating from the target as being incorrectly identified as the target. However, for metagenomes, such rates are as low as 0.0001, which seems small. However, when expressed as the number of reads per 10 million that were incorrectly identified as being the target (as in Figure 1), this is a more practical perspective. A false positive rate of 0.0001 translates to 1000 counts per 10 million, which is likely intolerable if the target were only be 100 counts per 10 million. The SEED and RefSeq databases required longer reads for more accurate matching of beta-galactosidase, preferably 200bp and above. Regardless of database choice, true positive rates increase with increasing read length (Figure 1A) but the number of false positives increase with increasing read length as well (Figure 1B). This results in increased sensitivity but decreased specificity with increasing read length (Figure S1). However, overall accuracy is increased with increasing read length (Figure S1).

**Figure 1.**
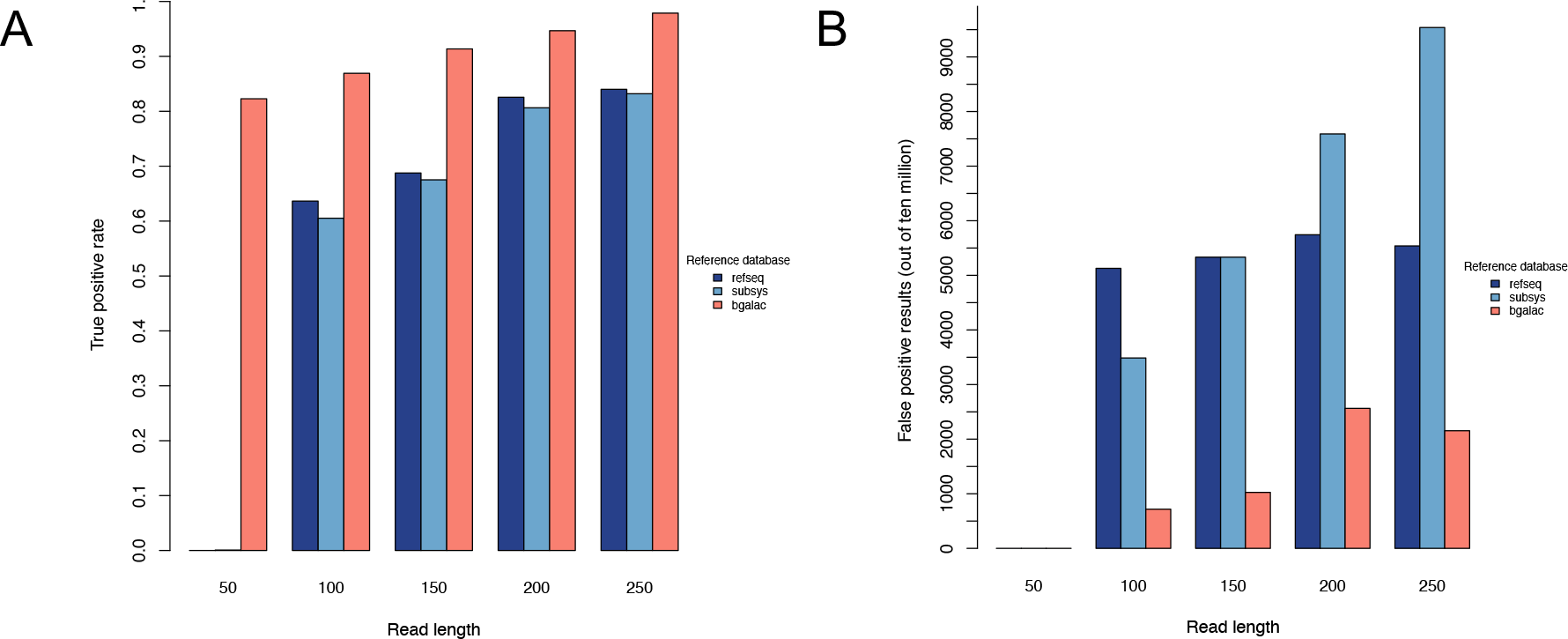
True (A) and false (B) positive rates, across different read lengths, of detecting the target beta-galactosidase sequence in simulated metagenomes using DIAMOND with “sensitive” flag and default e-value and 3 databases: NCBI RefSeq (navy), SEED Subsystems (blue), and beta-galactosidase (salmon).

### Selection of appropriate e-value threshold for mapping reads

To evaluate an appropriate DIAMOND e-value threshold for mapping reads of different lengths to a custom database, error rates were calculated for different e-values. We simulated a metagenome of 2,500 beta-galactosidase reads per 100,000 total reads with the non-beta-galactosidase reads being derived from other sequences in our known protein database (see Methods). Metagenomes were simulated at all 5 read lengths: 50, 100, 150, 200, and 250bp. Reads were aligned to the beta-galactosidase database using e-value cutoffs of 1e-3 (default DIAMOND e-value threshold), 1e-10, 1e-25, 1e-50, and 1e-100 (**Figure 2**). Both true and false positives decrease with more stringent e-values (Figure 2A-B), resulting in lower sensitivity but higher specificity (Figure S2). For every read length, the default e-value was most accurate for this unique situation where a custom database containing only the gene of interest is used to detect the gene of interest, beta-galactosidase, in the metagenome (Figure S2).

**Figure 2:**
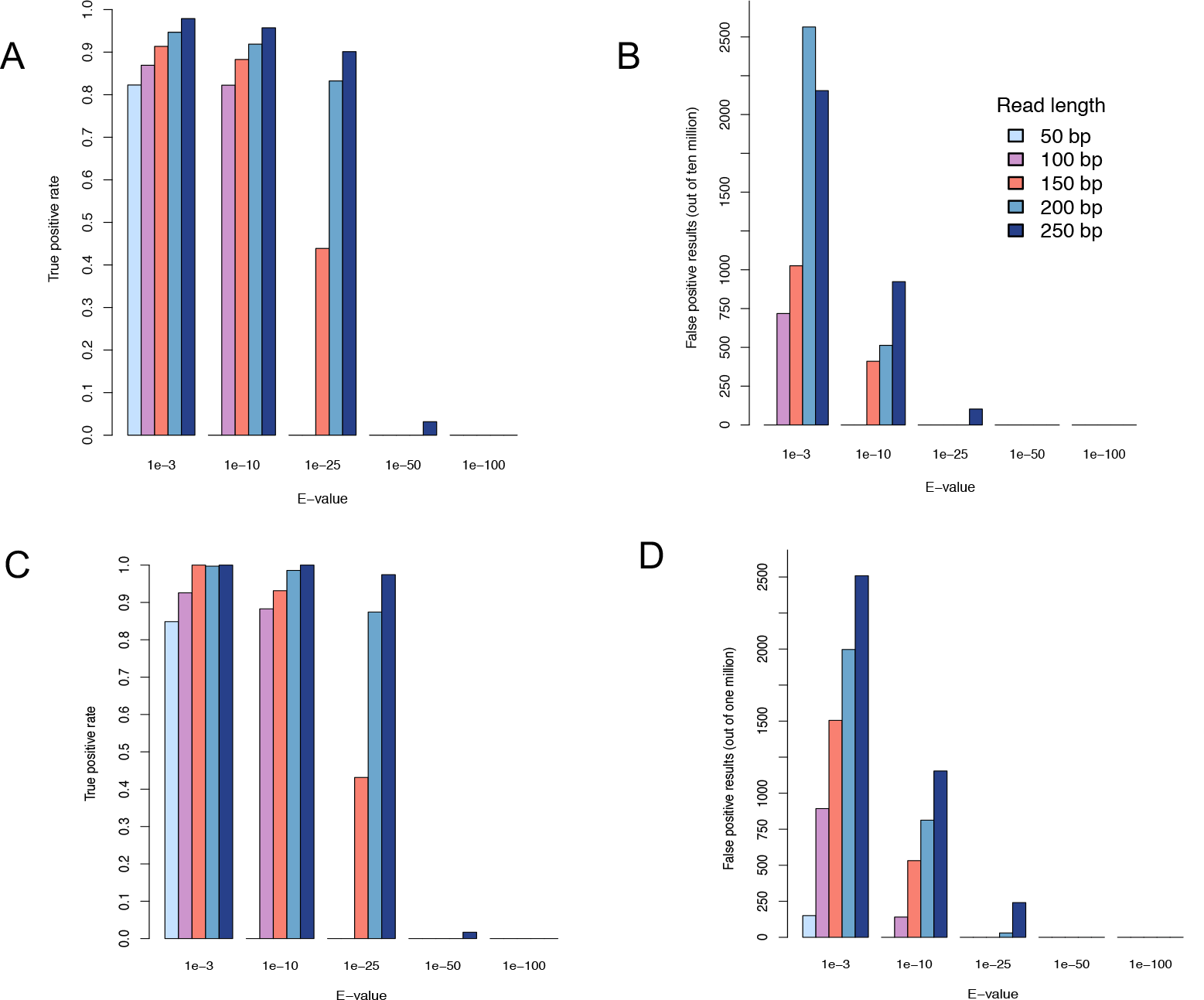
(A) True positive rates and (B) false positive rate results from simulated metagenomes containing 2,500 beta-galactosidase sequences at varying e-value cutoffs and read lengths of 50 bp (light blue), 100 bp (purple), 150 bp (salmon), 200 bp (blue), and 250 bp (navy). (C) True positive rates and (D) false positive rate results from simulated metagenomes containing 350 Homoserine O-acetyltransferase at varying e-value cutoffs and read lengths of 50 bp (light blue), 100 bp (purple), 150 bp (salmon), 200 bp (blue), and 250 bp (navy).

While the choice to examine beta-galactosidase in our previous experiments was due to our lab’s interest in this enzyme, we conducted another series of experiments with a protein sequence selected specifically to maximize the difficulty of distinguishing it from other proteins. Using our custom database of known proteins, we computed all pairwise percent identities of all sequences in the database and chose the protein with the highest percent identity with other proteins in the database. Homoserine O-acetyltransferase (HTA), which had 28 matches of mean 42% identity with other proteins in the database of known proteins, was selected for this experiment. Of the 28 sequences with high pairwise identity in the known protein database, only 14 are HTA and the other 14 proteins are not HTA. The 14 true HTA proteins were used to construct a custom database while all of the known proteins, including the 14 off-target sequences were used to create simulated metagenomes. Using simulated metagenomes containing 350 HTA reads per 100,000 total reads at all 5 read lengths, the metagenomes were aligned to a custom database containing the HTA protein sequences. To detect HTA, the best choice of e-value threshold was dependent on read length. For read lengths 100bp and higher, the best choice of e-value was not the default (Figure 2C-D). For 100-150bp, a threshold of 1-e10 yielded the highest accuracy (Figure S3). For 200-250bp read lengths, 1e-25 was most accurate (Figure S3). In general, the optimal e-value threshold increases with increasing read length.

### Choice of protein database

Protein databases differ in their organization, level of resolution, and level of evidence for annotations. To understand how the choice of protein database impacts the accuracy of functional annotation, several databases—Carbohydrate Active Enzyme (CAZy), NCBI nonredundant RefSeq, SEED Subsystems and a customized database containing only beta-galactosidases—were used to annotate metagenomes that were simulated from a database of proteins of experimentally verified function (see Methods). CAZy is a database of sequence-based families of enzymes that assemble, modify and breakdown oligo- and polysaccharides [12]. The NCBI RefSeq database contains a comprehensive set of non-redundant protein sequences [13]. The SEED subsystems database contains collections of functionally related protein families [14]. The customized database contained only beta-galatosidase sequences with previously verified functionality.

One hundred metagenomes of 100,000 reads were simulated from our database of known proteins with increasing doses of beta-galactosidase sequences ranging from 25 to 2,500 sequences for each of the five different read lengths: 50 bp, 100 bp, 150 bp, 200 bp, and 250 bp. Each metagenome was annotated against each database to determine the total number of beta-galactosidase hits. At all read lengths, the custom beta-galactosidase database was closer to the “expected” dose-response than the other databases (**Figure 3**). At 50bp, only the custom database performed well. At higher read lengths, the CAZy database appeared to over-estimate and the RefSeq and SEED subsystems databases appeared to underestimate the number of betagalactosidase reads present.

**Figure 3:**
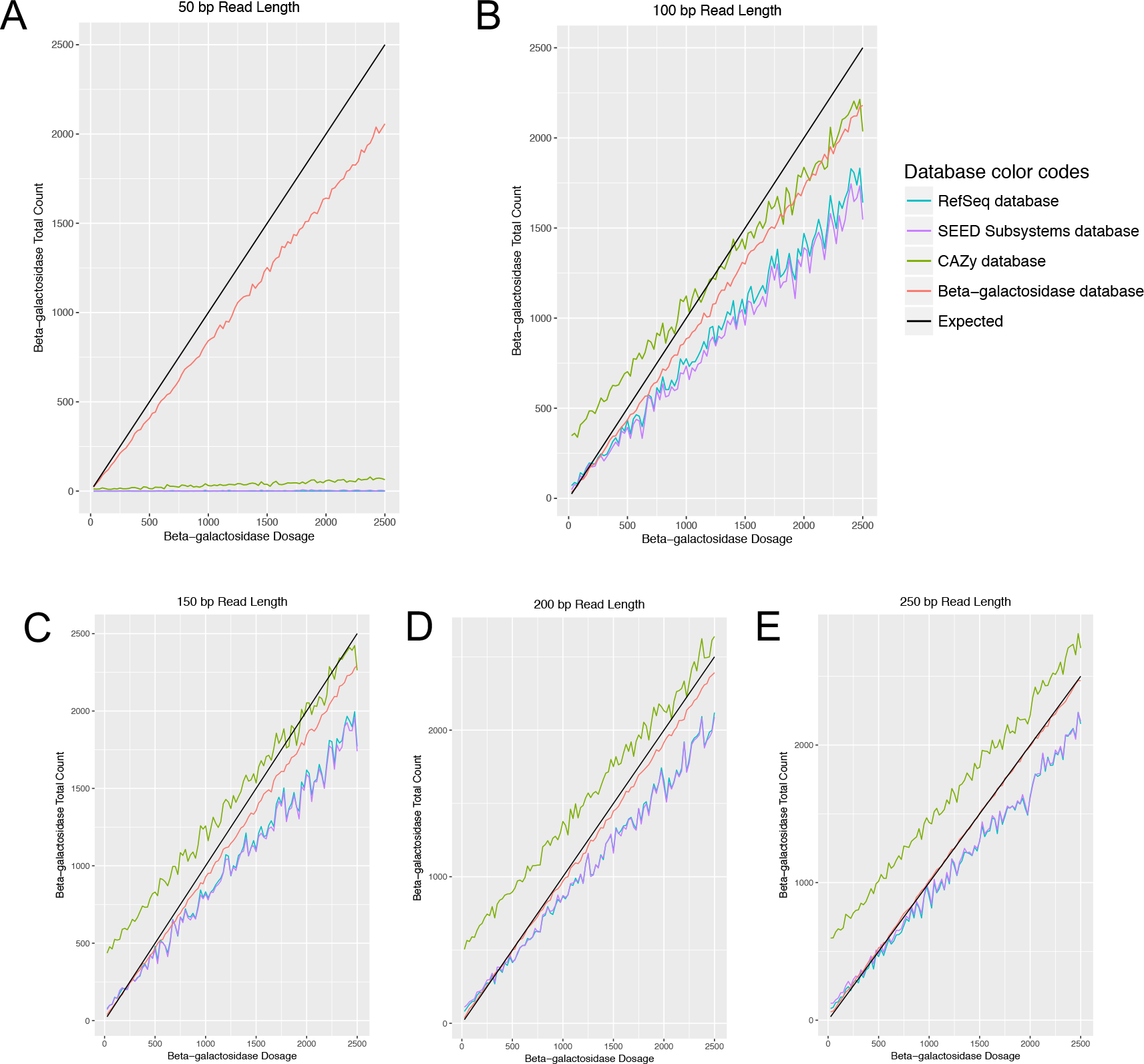
Comparison of total number of beta-galactosidase annotations across varying beta-gal dosages and read lengths of (A) 50 bp, (B) 100 bp, (C) 150 bp, (D) 200 bp, and (E) 250 bp between 4 databases: NCBI RefSeq (blue), SEED Subsystems (purple), CAZy (green), and Beta-galactosidase (red). The expected line is black.

To better understand why the CAZy database over-estimated beta-galactosidase abundance, we mapped the reads to CAZy families. For the beta-galactosidase dose experiment, some families showed an increasing dose while other families showed a constant number of hits across all doses (**Figure 4**). GH42 had the most obvious dose-response with a more moderate dose-response for GH2. Other GH families with potential beta-galactosidase members had an even response across the metagenomes. This phenomenon was true at all read lengths: 50bp, 100bp, 150bp, 200bp, and 250bp (Figure S4). Some of the CAZy families which contain beta-galactosidase (GH1, GH2, GH16, GH35, GH42, and GH98) also contain other genes that are not beta-galactosidase. This suggests that there may be off-target hits when using the CAZy database.

**Figure 4:**
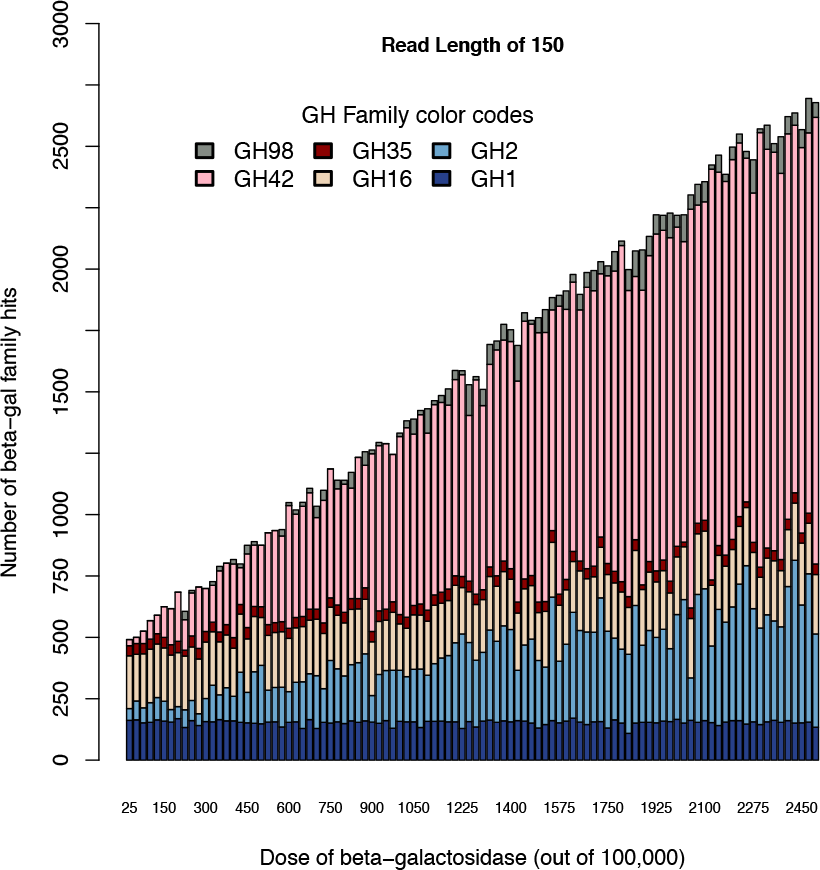
CAZy family hits with increasing beta-gal dosages at a read length of 150 bp. Annotations against 7 CAZy families that contain beta-galactosidase were included: GH1(navy), GH2 (blue), GH16 (tan), GH35 (red), GH42 (pink), and GH98 (grey).

### Evaluation of strategies for paired end reads

When mapping paired end reads to a protein database, mapping tools such as DIAMOND are not able to leverage the joint nature of pair-end reads. One must either map the two reads independently or the reads can be merged and then mapped. The extent to which sequences of the two ends can be merged depends on the size selection of the DNA prior to sequencing and the length of the reads. In the current study, we analyzed stool metagenomes from two projects with different read formats and insert sizes. The adult stool metagenomes were sequenced in a 2×101bp format from DNA with an insert size range of 280-320 bp [15]. With the larger insert size, the average percent of read pairs that could be merged in the 30 largest metagenomes used in the present study was 34.6% and the average length of the merged reads was 112 bp. The infant stool metagenomes were sequenced in a 2×151bp format from DNA with an average insert size range 119-289 bp [16]. In these metagenomes, which were deliberately sequenced with a smaller insert size to improve overlaps, we found that the average percent of read pairs that could be merged was 87.9% and the average length of the merged read was 188 bp. Thus, with a custom size selection step during library preparation, it is possibly to vastly improve the number of overlapping reads and the resulting length of the merged reads.

We next evaluated whether an alternative strategy should be used when analyzing shotgun metagenomes in the paired end format with few overlapping paired reads. Other options besides merging reads included using only one of the read pairs (referred to here as the “R1 strategy”) or independently mapping both reads and keeping only those hits in which both reads map to the same protein (referred to here as the “congruent strategy”). We used our custom database for beta-galactosidase, a well-known enzyme present in both infant and adult stool, and mapped reads against this beta-gal database using the R1, congruent, or merged read strategy. With the infant metagenomes, in which most reads were mergable and the resulting reads were not much longer than the single reads, both alternative strategies were highly correlated with the merged strategy, as expected. The correlation between merged and R1 strategies was r=0.97, p<2.2e-16 (**Figure 5A**) and the correlation between merged and congruent strategies was r=0.96, p<2.2e-16 (**Figure 5B**). Furthermore, the number of reads mapped using any of the strategies was similar, likely due to the high percent of overlapping reads (88%). With the adult metagenomes, in which only one third of the reads overlapped, the alternative strategies had a lower correlation with the merged strategy: the correlation between merged and R1 strategies was r= 0.91, p<4.9e-12 (**Figure 5C**) and the correlation between merged and congruent strategies was r= 0.95, p<1.1e-14 (**Figure 5D**). Roughly double the number of counts were obtained with the R1 strategy compared with the merged strategy. The fewest counts were obtained with the congruent strategy, although the correlation was higher. In summary, it is best to increase the number of overlapping reads during the library preparation phase; however, if few reads can be merged, the congruent strategy appears to be a viable alternative.

**Figure 5:**
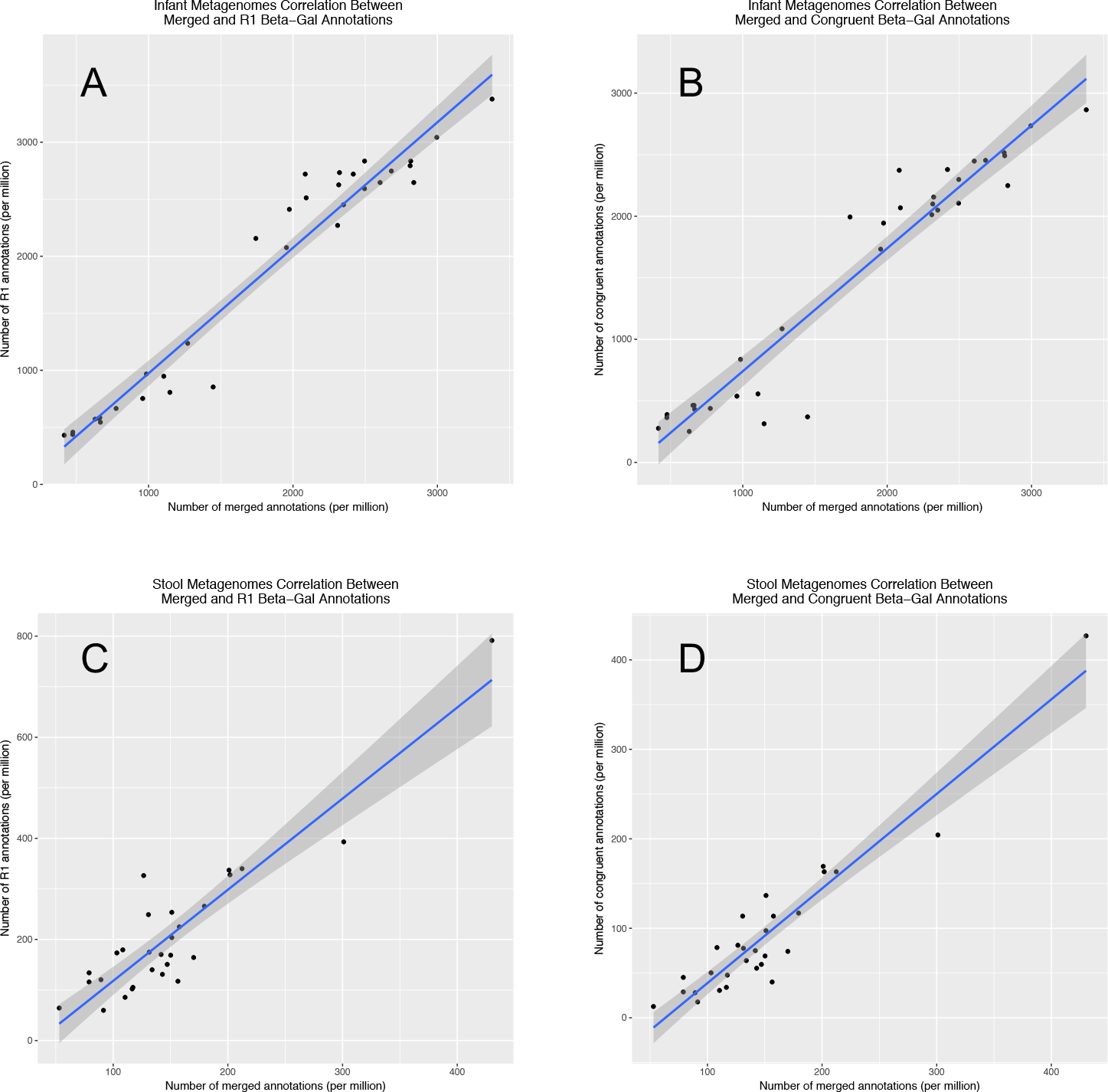
Correlations of the R1, congruent, and merged read annotations in (A-B) infant fecal metagenomes 2×150bp with nearly 90% merge-able reads and (C-D) adult fecal metagenomes 2×100bp format with only one third merge-able reads.

### Evaluation of the effect of sequencing depth

The choice of sequencing depth would be expected to effect the likelihood of detecting all the proteins in a given population. To determine the minimum sequencing depth needed for metagenomic sequencing of stool metagenomes, we sub-sampled stool metagenomes that were deeply sequenced. The 33 infant metagenomes had a range of sequencing depths from 13.7 to million reads per metagenome [16]. We also selected the 30 largest metagenomes from among 500 metagenomes in an adult stool metagenome project [15]. Each metagenome was randomly sub-sampled to produce metagenomes ranging from 100,000 to 10 million sequences each, with 10 sub-samples at each read depth. The standard deviation of the counts of the target gene, beta-galactosidase, across the 10 sub-samples was then calculated at each read depth. As expected, the greater the sequencing depth in infant stool metagenomes, the lower the standard deviation relative to quantitation from the full metagenome (**Figure 6**). For quantification of beta-galactosidase in infant metagenomes, it appears that even a few million merged reads is sufficient to accurately capture the relative abundance of beta-galactosidase. We repeated the experiment with adult stool metagenomes but because these were prepared with suboptimal insert sizes for merging, even the largest metagenomes had only 4 million merged reads, which would be insufficient for a sub-sampling experiment. We therefore applied the conguent strategy to all unmerged read pairs in the largest metagenomes with at least 20 million reads. The standard deviation of the count of beta galactosidases in the sub-sampled metagenomes relative to the full metagenomes reached a plateau near 10 million reads (Figure 6B).

**Figure 6:**
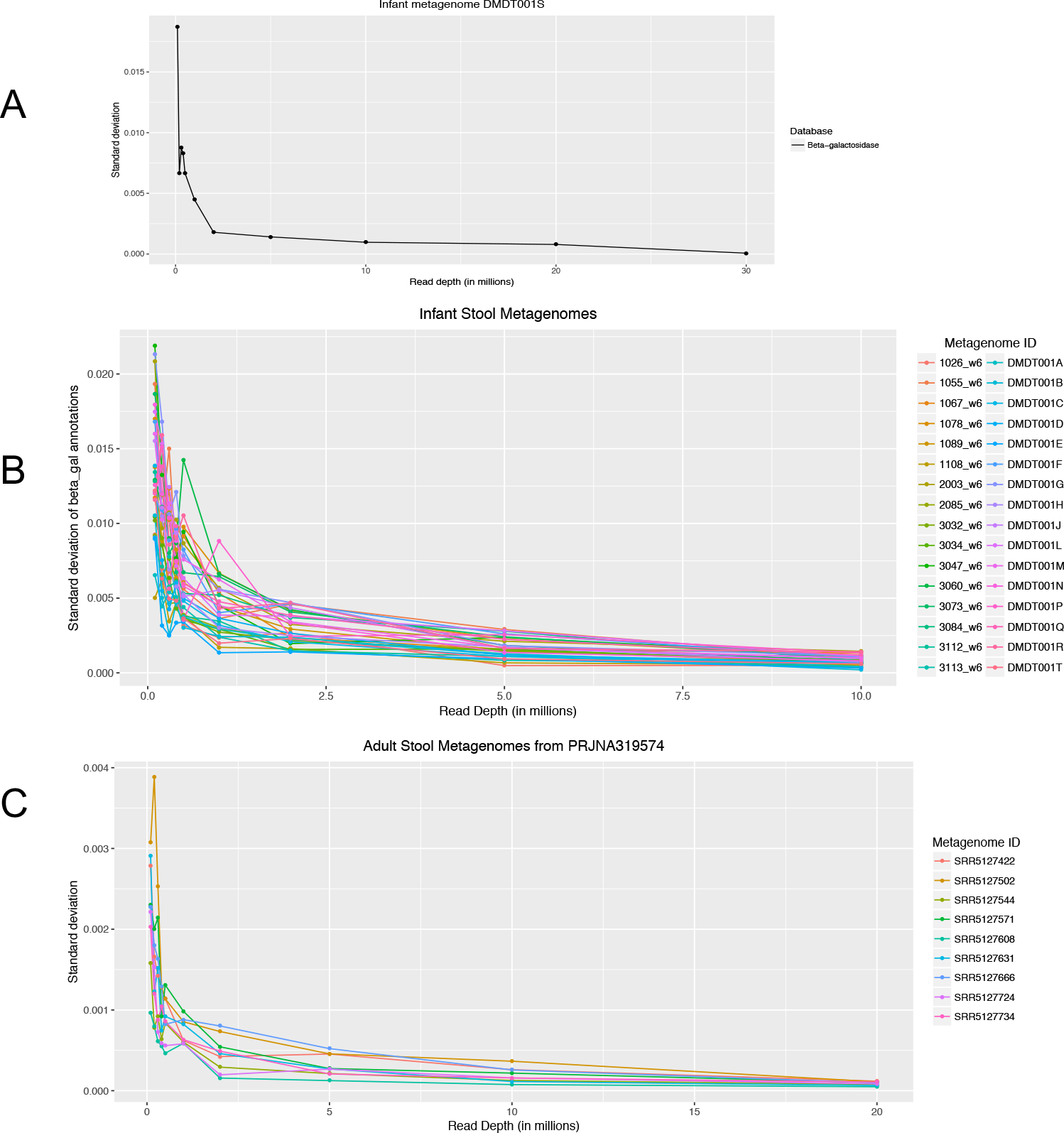
Effect of sub-sampled read depth on the standard deviation of target gene hits in the sub-sampled metagenome relative to the full metagenome. The target gene was beta-galactosidase, mapped using a custom database of beta-galactosidase sequences. The metagenomes tested were (A) an infant fecal metagenome of 30 million merged reads, (B) all infant fecal metagenomes of at least 10 million merged reads, and (C) adult fecal metagenomes with at least 20 million unmerged reads mapped using the congruent strategy.

Given that one of the infant metagenomes was very deeply sequenced (30 million reads with merging of nearly 90% of reads), we conducted further experiments to determine the read depth to classify other groups or enzymes of interest. For CAZy families GH29 or GH95 which contain fucosidases, 5 million reads appear to be sufficient for quantification (Figure S5). This appears to be similarly true for broader classifications of “animal carbohydrates”, “plant cell wall carbohydrates”, “mucin”, and “sucrose/fructans” based on multiple CAZy families (**Figure 7A**).

**Figure 7:**
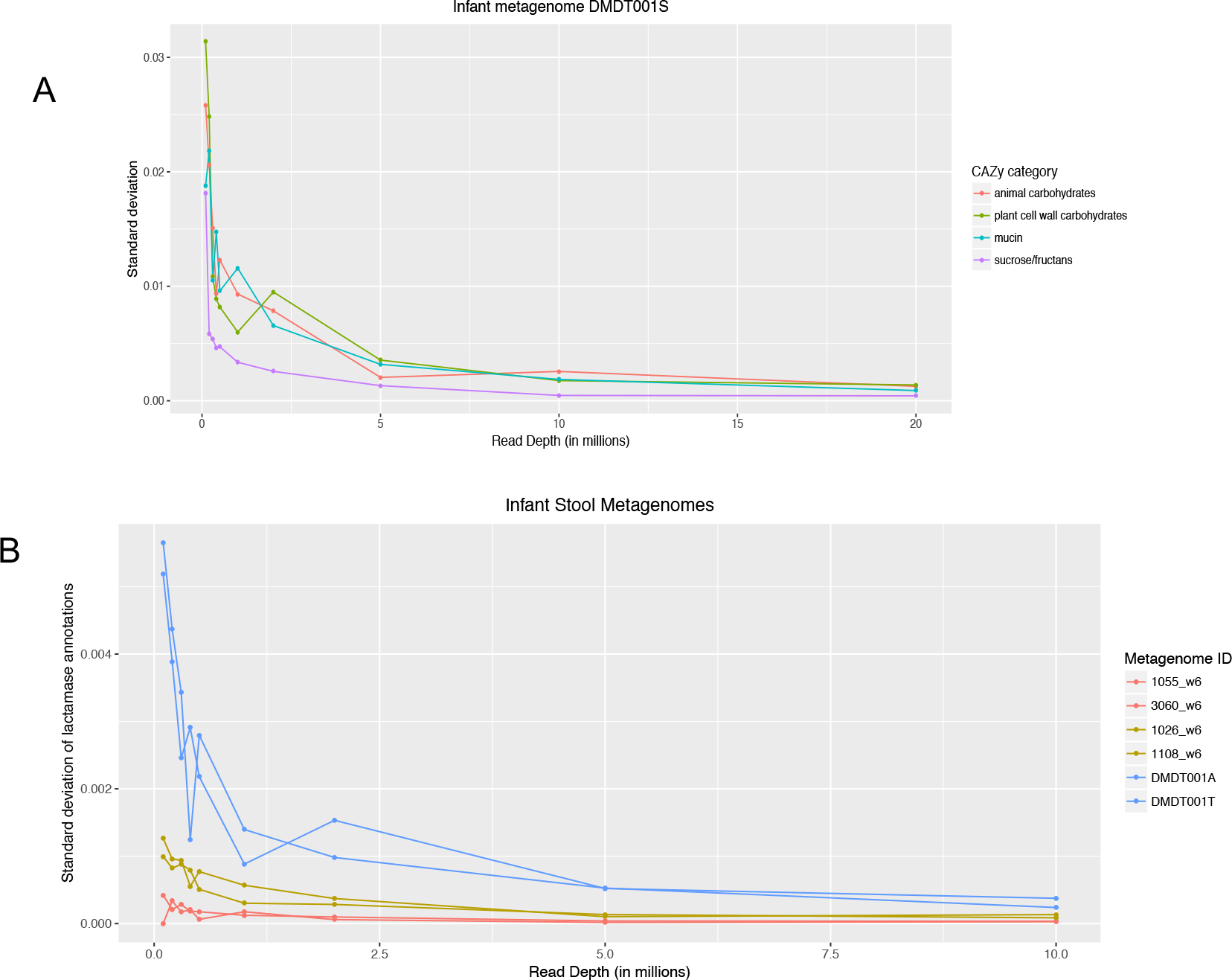
Effect of sub-sampled read depth on the standard deviation of target gene hits in the sub-sampled metagenome relative to the full metagenome. The target gene was (A) CAZy category, mapped using the CAZy database, or (B) beta-lactamase, using a custom database. The metagenomes tested were (A) an infant fecal metagenome of 30 million merged reads, and (B) six infant stool metagenomes representative of high (blue), medium (yellow), and low abundance (red) in beta-lactamase.

Antimicrobial resistance genes in fecal metagenomes are of practical importance, but they are rare. We therefore asked what the sequencing depth would need to be to accurately quantitate an antimicrobial resistance gene, beta-lactamase, that was known to be both low abundance and differentially abundant in the infant metagenomes [16]. We sub-sampled six infant genomes—two known to be high in beta-lactamase, two with medium beta-lactamase and two with low beta-lactamase—and mapped against a custom database for beta-lactamases. The standard deviation of beta-lactamase counts, relative to the full metagenomes, seemed to be again minimized by 5 million merged reads (**Figure 7B**). In general, 5 million merged reads seem sufficient for functional assessment of infant stool metagenomes.

## Discussion

As the cost of sequencing continues to decrease, more studies will use shotgun metagenomes to estimate the functional content of fecal microbiomes and scientists will be faced with practical choices in the pre- and post-sequencing phases. In the current study, we found that read length, e-value threshold, choice of protein database, strategy for paired end reads, and sequencing depth all impacted the ability to detect sequences of known abundance in shotgun metagenomes of human fecal microbiomes. Furthermore, decision-making around these parameters is intertwined as they affect each other.

Previous studies have observed a relationship between read length and the accuracy of gene detection [11, 17]. Consistent with their observations, we found that overall accuracy increases with read length. What is more interesting is translating this knowledge to which of the commonly available sequencing formats are therefore appropriate. SR50 should clearly never be used; SR100 should only be used with a small customized database, provided that quality trimming does not extensively trim the reads. PE100 should only be used for the same purpose as SR100 or it can be used more broadly if the sequencing library is size selected to enable overlapping paired ends, which are then merged to create merged reads longer than 100bp. When processing longer read formats, reads <100bp should simply be discarded. The Human Microbiome Project’s WGS Read Processing protocol retaining all reads >60bp [18]; clearly, this threshold is too low for functional annotation of gene sequences. Given today’s read format choices, the ideal choice would be PE150 with size-selection to enable overlapping reads. This can generate merged reads with a median length near 250bp (D. Lemay, unpublished observations).

We also investigated what to do with legacy data in which most paired end reads are not mergable. For one of the largest sets of fecal metagenomes from healthy adults to date [15], we found that only one third of the reads overlapped and when they did, the mean read length was still rather short (just 112bp). In another dataset [16], merging reads was much more successful ((~88% overlap, mean 188bp, on PE150 data). Using these two datasets, we compared three read strategies: a “merged strategy” (using only merge-able reads), an “R1 strategy” (using only forward reads), and a “congruent strategy” (independentally mapping both reads and keeping only those hits in which both reads map to the same protein). As expected, all three strategies are highly correlated with “merged” and “congruent” having the highest correlation. Therefore, the “congruent” strategy is a reasonable alternative when reads are not mergeable.

For the design of future experiments, should the sequence library be size-selected to enable overlapping pairs of reads? If functional annotation is a primary focus, then merged reads are highly desirable. Fewer hits were obtained with the “congruent” strategy, suggesting that the sensitivity would be lower if paired end reads cannot be merged. Rare genes, such as antimicrobial resistance genes, would then be more difficult to quantitate.

Knowing that appropriate e-value thresholds are dependent on the search tool used as well as read length and database size [17], we investigated what thresholds would be appropriate for use with DIAMOND at different read lengths. We found that two different protein sequences had two different optimal thresholds for accurate detection. This supports the opinion of other researchers that it may not be possible to identify a single cut-off for all proteins [7, 17]. However, given that we identified and tested a “worse case scenario”, our data suggests that it is possible to identify reasonable ranges for mean read lengths. For example, using DIAMOND in “sensitive” mode, a threshold within the range of the default 1-e3 to 1-e10 would be appropriate for read lengths of 100-150bp when a small customized database is used. A threshold within the range of 1e-10 to 1e-25 would be appropriate for read lengths of 200-250bp. For a larger database, like the SEED, we observed higher false positives with increasing read length, suggesting that a threshold more strict than the default is needed for broad databases. Generally, if the research question in mind has low tolerance for false positives, thresholds towards the stricter end of the suggested ranges should be chosen.

As it is well-known that shotgun metagenomes will have reads of varying length, it may be useful to implement a binning strategy to apply e-value thresholds proportional to the read length. First, shorter reads (<100bp) should be removed. Then, if the read lengths vary dramatically within a sample, it may be useful to sort each sequence or merged pair of sequences into bins such that appropriate e-value thresholds can be applied to each bin.

Bengtsson-Palme has suggested that a database that is specialized for the research question should be used and, if it does not exist, should be constructed based on genes of verified function described in the literature [7]. Our data supports that opinion. With the goal of detecting beta-galactosidase, the custom database for this enzyme was far superior to the other databases. NCBI and SEED tended to under-estimate the number of beta-galactosidase enyzmes in the simulated metagenome, while the use of CAZy over-estimated them. The fact that beta-galactosidases exist in several distinct families in the CAZy database and that those families, in turn, include some sequences that are likely not beta-galactosidases further demonstrates the difficulty interpreting results for specific enzymes using the CAZy database. Criteria previously suggested for customization of databases include whether the sequences are experimentally verified, the quality of the data, and the functional and/or taxonomic coverage of the data [19]. If analyzing the content of a particular enzyme of interest, the database could be focused on experimentally verified sequences of that enzyme as we did for beta-galactosidase. By extension, if a particular pathway is the focus of the hypothesis, the protein sequences associated with pathway should be curated for the database. Hypothesis-specific databases would be faster, more accurate, and highly interpretable.

Investigating the effect of sequencing depth (e.g. library size) on the ability to quantitate specific genes or functional categories in human fecal metagenomes, we found that 5 million merged reads or 10 million unmerged reads would be sufficient to quantitate even rare genes. In prior work, Nayfach and Pollard surprisingly found that reducing the sequencing depth of human gut metagenomes by 95% introduced < 2.5% variation in gene copy number estimates [20]. However, they also suggest that the effects differ for common versus rare genes. What is rare? Considering that there are, on average, around 500,000 unique microbial genes per individual’s gut microbiome [21], many individual genes will be rare. We found that beta-galactosidase, the enyzme that digests lactose, has about 500 copies per 5 million merged reads in an adult metagenome and tens times as many in an infant metagenome. For beta-lactamase, an antimicrobial resistance gene, there were about around 100 copies per 5 million merged reads in a deeply sequenced infant metagenome. The concept of shallow sequencing—roughly 500,000 sequences per metagenome—has been put forward as an in-between alternative between 16S sequencing and current shotgun metagenomes [22]. With 500,000 unique microbial genes per individual’s gut microbiome [21], shallow sequencing would sample each gene only once, on average, which would not be sufficient to assign presence/absence to most genes with confidence, nevermind comparative quantitation among samples. Answering hypotheses about specific genes or functions more narrow than say, “carbohydrate metabolism”, will require a more deeply sequenced metagenome. We suggest a minimum of 5 million merged reads of at least 150bp in length.

Our recommendations are limited to shotgun sequencing of human fecal samples. These samples are high biomass with nearly all DNA from microbial sources and low risk of substantial contamination. Furthermore, host DNA is easly removed [23, 24]. Others have indicated that problem of higher variation due to contamination in low biomass samples [25] which suggests that recommendations based on human fecal samples may not extend to other samples. It is possible that recommendations may not even extend to stool samples of other animals. Zaheer et al reported that >50 million reads per sample would be needed to characterize the fecal resistome of cattle fecal samples [26]. Nearly all reads (>97%) were uncharacterizable in that study; the authors suggested that this may be due to the presence of feed-associated plants that were not in the database.

The experiments in the current study do not address how to use gene abundances to compare across metagenomes. Nayfach and Pollard review three methods: (1) gene relative abundance, which is relative amounts of genes found in a sample, (2) average genomic copy number, which is the expected number of copies of the gene per cell, and (3) gene absolute abundance, which cannot be estimated from sequence data alone [20]. The choice should probably be dependent upon the application. If the primary goal is to calculate the carbohydrate active enzyme profile of a fecal metagenome without regards to whether those enyzmes are derived from bacteria with a reference genome or even non-bacteria, such as archaea or yeasts, then the gene relative abundance may be expected to yield less bias than average genomic copy number. However, it is important that differential abundance testing is done using a count-adapted method, such as DESeq2 [27] and edgeR [28], which simultaneously account for differences in library sizes and the compositional nature of the data and have demonstrated applicability for metagenome data [29]. A comparison of 14 different methods demonstrated that these methods have the best performance to determine differnentially abundant genes in metagenomes [30].

Our study is intended to be agnostic to the pipeline used, except for the use of DIAMOND as a mapping utility. An argument against using point-and-click computational computational pipelines is the inability to assign appropriate cut-offs for genes of interest [7]. An advantage of creating one’s own pipeline or modifying one that is open source, such as SAMSA2 [31], is the ability for users to define e-value thresholds and use customized databases. SAMSA2, although built for metatranscriptomes, also works to assess gene abundances in metagenomes with the omission of the rRNA removal step, which is unnecessary for metagenomes. Therfore, a version of the SAMSA2 pipeline with that modification was used for the experiments in this paper. For the functional annotation of fecal metagenomes, it could be argued that a broad overview with mappings to the Clusters of Orthologous Genes (COG) database [32], KEGG Orthology [33], and/or SEED subystems [14] databases is a reasonable first step followed by tests for specific hypotheses with databases customized for those hypotheses.

An alternative to the DIAMOND/BLAST approach would be a mapping strategy which uses hidden Markov models (HMMs). A profile HMM encodes the statistical probability of each amino acid in the sequence of a protein family used to train the model. This can be useful to identify distant homologs that may not be represented in a database. The Pfam database is a large collection of protein families, with each family represented by a multiple sequence alignment and HMM [34]. Pfam also collects entries into clans that are collections of proteins related by profile HMM. Several tools, such as HMM-GRASPx [35] and MetaCLADE [36], identify protein domains in metagenome and/or metatranscriptome sequences. These methods are likely most useful to annotate sequences when proteins are not well-represented in reference databases, such as in environmental sequence data [37], because protein domain information is better than no information. However, proteins with very different functions can share the same protein domain and there can be multiple protein domains in the same protein. For human fecal metagenomes, it is now possible to map 70-90% of reads to genes [38] using the integrated gene catalog (IGC) of the human gut metagenome [39]. Given that the average percentage of prokyarotic genomes that comprise gene-coding regions is 87%, mapability of reads from human gut microbiomes has approached the maximum achievable [39]. Thus, it is both possible and preferable to identify the complete protein rather than a protein domain to enable interpretation of the results. In theory, a profile HMM can be built using complete sequences from different taxa for a particular protein; whether such a strategy has higher performance than the DIAMOND/BLAST approach is beyond the scope of the current analysis.

Assignment of sequence reads from fecal metagenomes to broad functional categories gives the impression of stability [40, 41], but this impression is likely false when investigating function on a finer scale of individual genes. Publicly available human fecal metagenomes have been reanalyzed for functional content using alternate methods [42] or for meta-analyses across cohorts [38]. Legacy data should also be re-analyzed for specific hypotheses. The results of the current study suggests some guidelines for using legacy data to address hypotheses of interest: create a custom database of the gene or genes of interest, merge paired ends to create longer reads, and adjust the e-value threshold for the median read length or bin reads by read length for different e-value thresholds.

## Conclusions

The accurate identification of sequences of known abundance in human fecal metagenomes was affected by read length, e-value threshold, choice of protein database, strategy for paired end reads, and sequencing depth. If the primary purpose of metagenomics analysis is to infer specific functions, then DNA extracted from human fecal samples sequenced using the Illumina platform should be size-selected to enable merging of paired end reads, and should be sequenced in the PE150 format or better with a minimum sequencing depth of 5 million merge-able reads, which will likely be closer to 10 million reads. Expecting the merged reads to be 180-250bp in length, the appropriate e-value threshold for DIAMOND would then be more strict than the default. Accurate and interpretable results for specific hypotheses will be best obtained using small databases customized for the research question.

## Methods

### Construction of a database of proteins with known function

In order to test various functional metagenomic techniques, a database containing only proteins with experimentally verified functions was constructed. The three sources for protein sequences were Swiss-Prot [43], New England BioLabs (NEB) Inc., and those previously experimentally verified in *Bifidobacterium* [16–24]. From Swiss-Prot, a fasta formatted database of reviewed bacterial sequences with experimentally verified functions was downloaded. Sequences of experimentally validated glycosidases were sent directly from the NEB techical team. We manually collected a list of *Bifidobacterium* enzymes that were biochemically confirmed for their function [16–24]. The protein database (~5000 protein sequences) was then blasted against itself to remove all sequences with greater than 50% identity using custom scripts, removing ~1000 protein sequences that were closely related to other sequences within the database. Lastly, we reverse translated the protein database into a nucleic acid database using the *E. Coli* codon table and the tool EMBOSS Backtranseq [53]. Note, because DIAMOND translates nucleic acid sequences into protein sequences, the choice of codon table did not make a difference in this scenario.

### Construction of simulated metagenomes with increasing dosage of beta galactosidase

The database of proteins with known functions was split into two databases, one with only beta-galactosidase sequences and the other containing all other protein sequences. Sequences were randomly selected from the two databases to create 100 test databases with an increasing proportion of beta-galactosidase enzyme sequences (1/4,000 to 100/4,000). Next, *in silico* metagenomes were created using the next generation sequencing simulator software MetaSim [54]. Using MetaSim, 100 metagenomes of 100,000 reads with increasing dosages of beta-galactosidase (25/100,000 - 2,500/100,000) were simulated at 5 different read length of 50 bp, 100 bp, 150 bp, 200 bp, and 250 bp. Using the same reverse translated nucleic acid database, each sequence was extended by 20% (10% to each end) by randomly adding nucleotides based on actual nucleotide frequencies to represent more realistic DNA fragments with flanking intergenic regions. Another 500 metagenomes were simulated from the elongated sequences of increasing beta-galactosidase proportions at the 5 different read lengths.

### Fecal metagenome datasets

Two human fecal shotgun metagenomic datasets were analyzed in this paper. The dataset of infant fecal metagenomes included 33 samples [16]. The second dataset included 30 adult fecal metagenomes which were among the largest metagenomes from a cohort of 471 healthy, Western European adults of at least 18 years of age, as part of the 500 Functional Genomics project [15]. The infant metagenomes’ sequence data are in DDBJ Center with SRA accession no. SRP133760, and the adult metagenomes’ sequence data are in NCBI SRA with accession no. PRJNA319574.

### Metagenomic sequence analysis

The SAMSA2 pipeline [55] was modified for metagenomics analysis with the following steps: 1) reads mapping to the human genome were first removed with BMTagger [23], 2) paired-end reads were merged using PEAR [56], 3) sequence adaptor contamination and low quality bases were removed using Trimmomatic [57] and 4) the quality reads were then annotated against a protein reference database using DIAMOND, a high-throughput squence aligner [10]. DIAMOND is highly sensitive and runs at a speed that is up to 20,000 times faster than BLASTX and up to 2,500 times faster with the “sensitive” option. The pipeline script, master_beta.galac.db_analysis_stoolmg.sh, is in https://github.com/mltreiber/functional_metagenomics To determine gene content, reads were mapped against three publicly available databases, NCBI RefSeq [58], SEED Subsystems [14], and CAZy [12] with only the best hit retained. Reads were also mapped against a custom-built database which was a fasta-formatted list of experimentally verified beta-galactosidase sequences.

### Metrics of evaluation

The performance of sequence read mapping was evaluated using several metrics. Sequence reads originating from the target that were also classified as the target were considered true positives (TP). Sequence reads not originating from the target that were classified as the target were considered to be false positives (FP). Sensitivity refers to the ability to detect the target sequence while specificity refers to the ability to correctly identify sequences that are not the target; accuracy is the proportion of true results. Sensitivity, specificity, and accuracy were calculated as follows:

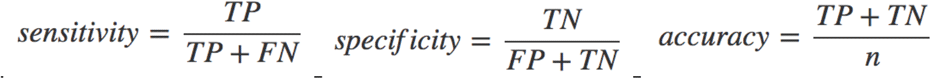

False negatives (FN) were target sequence reads that were not correctly classified as the target. True negatives (TN) were non-target sequence reads that were correctly classified as non-target. The number of total reads was equal to *n*.

## List of abbreviations

DIAMOND: a BLAST-like sequence aligner that uses translated sequence queries to greatly increase annotation speed over traditional BLAST.
MG-RAST: the MetaGenomics Rapid Annotation using Subsystems Technology server, a public analysis pipeline for handling metagenome and metatranscriptome datasets
PE: paired end read
SAMSA2: Simple Analysis of Metatranscriptomes through Sequence Annotation version 2, a standalone metatranscriptomics analysis pipeline
SEED: a protein database that seeks to group sequences into hierarchical categories, created by the Fellowship for Interpretation of Genomes (FIG group)
SR: single read

## Declarations

### Ethics and consent

Not applicable. The current study used publicly available data.

### Consent for publication

Not applicable.

### Availability of data and material

Human fecal metagenomes analyzed in the current study are available at DDBJ Center with SRA accession no. SRP133760 and in NCBI SRA with accession no. PRJNA319574. The database of experimentally verified protein sequences developed for this study, as well as other databases used, is in the databases directory of https://github.com/mltreiber/functional_metagenomics. The SAMSA2 analysis pipeline, modified for metagenomics, is called “master_beta.galac.db_analysis_stoolmg.sh”, and is available in the scripts directory of https://github.com/mltreiber/functional_metagenomics.

### Authors’ contributions

DGL conceived of the study. MLT conducted all experiments. MLT, IK, and DGL contributed to code. All authors interpreted data. MLT and DGL drafted the manuscript. DT and IK contributed to the manuscript. All authors read and approved the final manuscript.

## Acknowledgements

The authors thank Dr. Nina Kirmiz for the collection of *Bifidobacterium* enyzme sequences and Shannon E.K. Joslin for testing removal of human reads with BMTagger. The authors also thank Dr. Jinxin Liu, Dr. Lutz Froenicke, Dr. Matthew Settles, Dr. Zhengyao Xue, and Dr. Mary Kable for helpful discussions or comments.

## Additional files

**Additional file 1 – Supplementary Figures**

This PDF file contains supplementary figures, Figures S1-S5.

