## Supplemental Figures for "Pre- and post-sequencing recommendations for functional annotation of human fecal metagenomes"

Figure S1

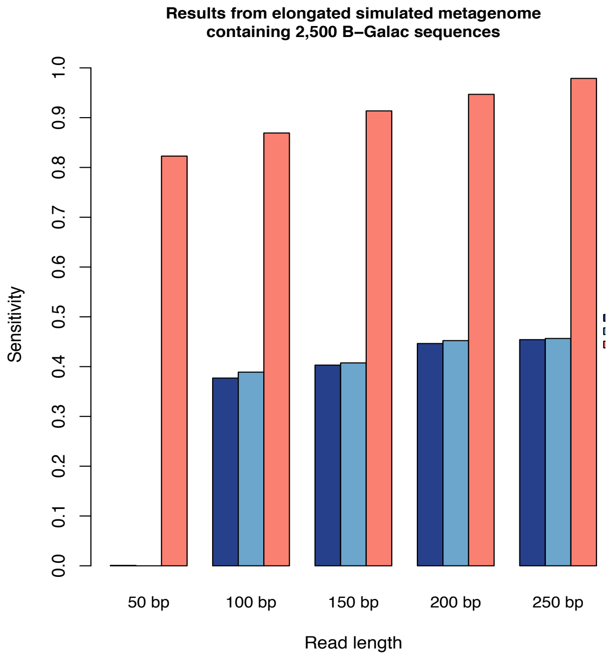


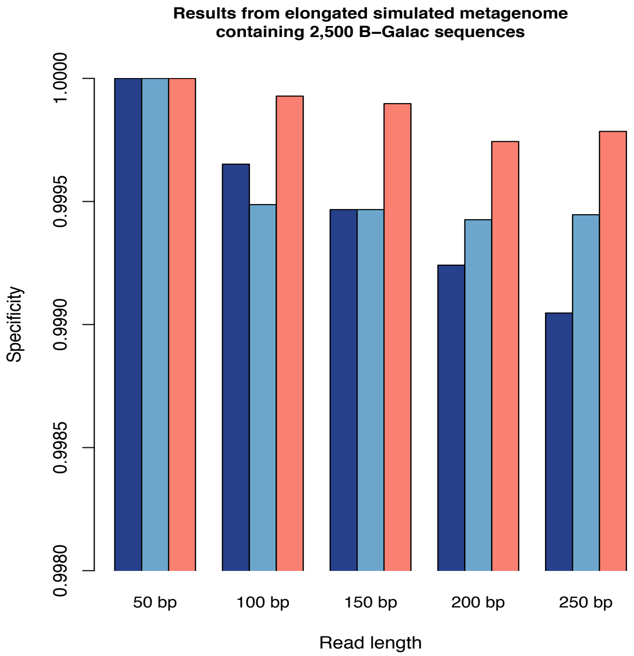


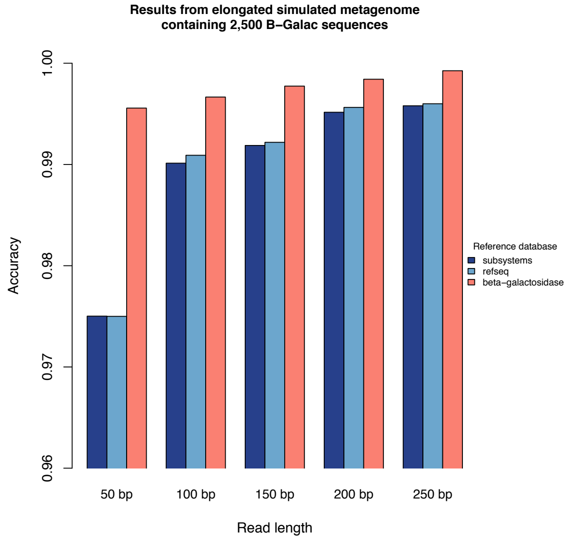


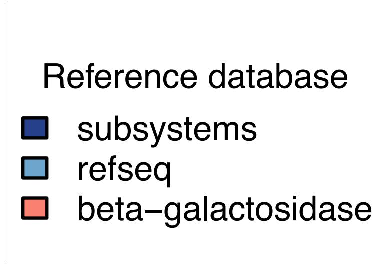


**Figure S1.** Sensitivity, specificity, and accuracy across different read lengths, of detecting the target beta-galactosidase sequence in simulated metagenomes using DIAMOND with "sensitive" flag and default e-value and 3 databases: NCBI RefSeq (navy), SEED Subsystems (blue), and beta-galactosidase (salmon).

### Figure S2

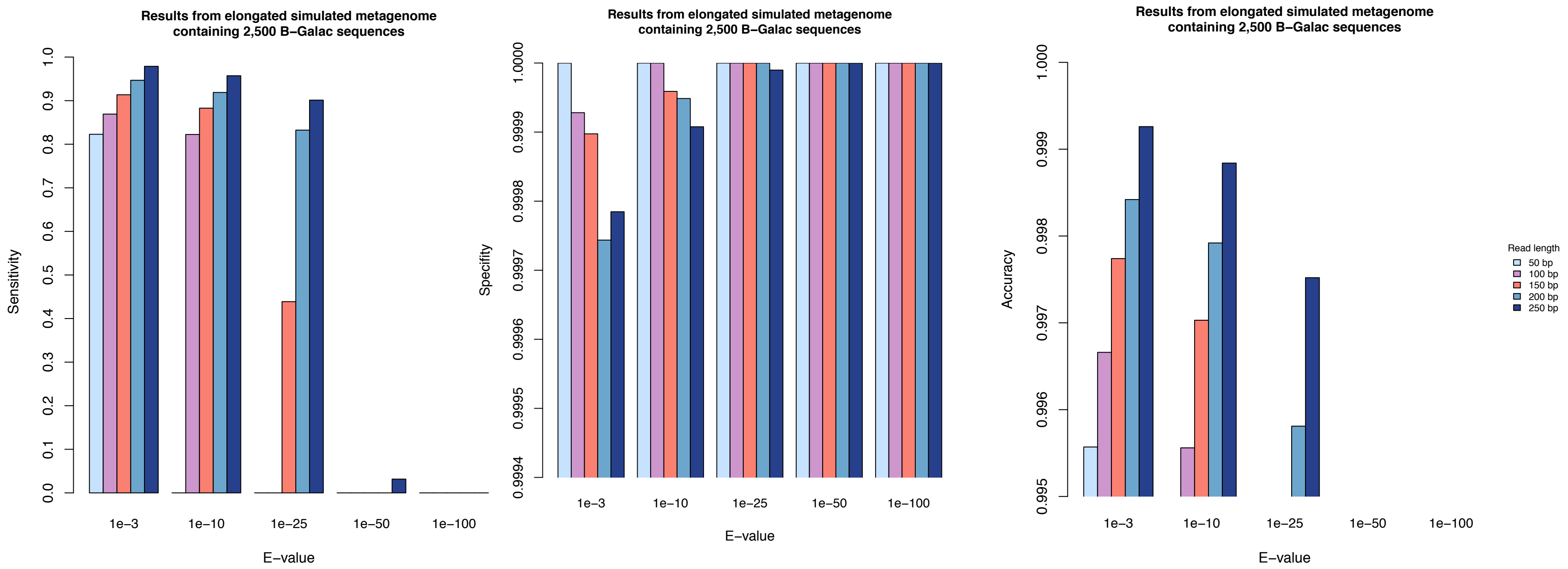
Results from elongated simulated metagenome containing 2,500 B−Galac sequences
Sensitivity
0.0
0.1
0.2
0.3
0.4
0.5
0.6
0.7
0.8
0.9
1.0
1e−3
1e−10
1e−25
1e−50
1e−100
E−value
Results from elongated simulated metagenome containing 2,500 B−Galac sequences
Specifity
0.9994
0.9995
0.9996
0.9997
0.9998
0.9999
1.0000
1e−3
1e−10
1e−25
1e−50
1e−100
E−value
Results from elongated simulated metagenome containing 2,500 B−Galac sequences
Accuracy
0.995
0.996
0.997
0.998
0.999
1.000
1e−3
1e−10
1e−25
1e−50
1e−100
E−value
Read length
50 bp
100 bp
150 bp
200 bp
250 bp

### Figure S3

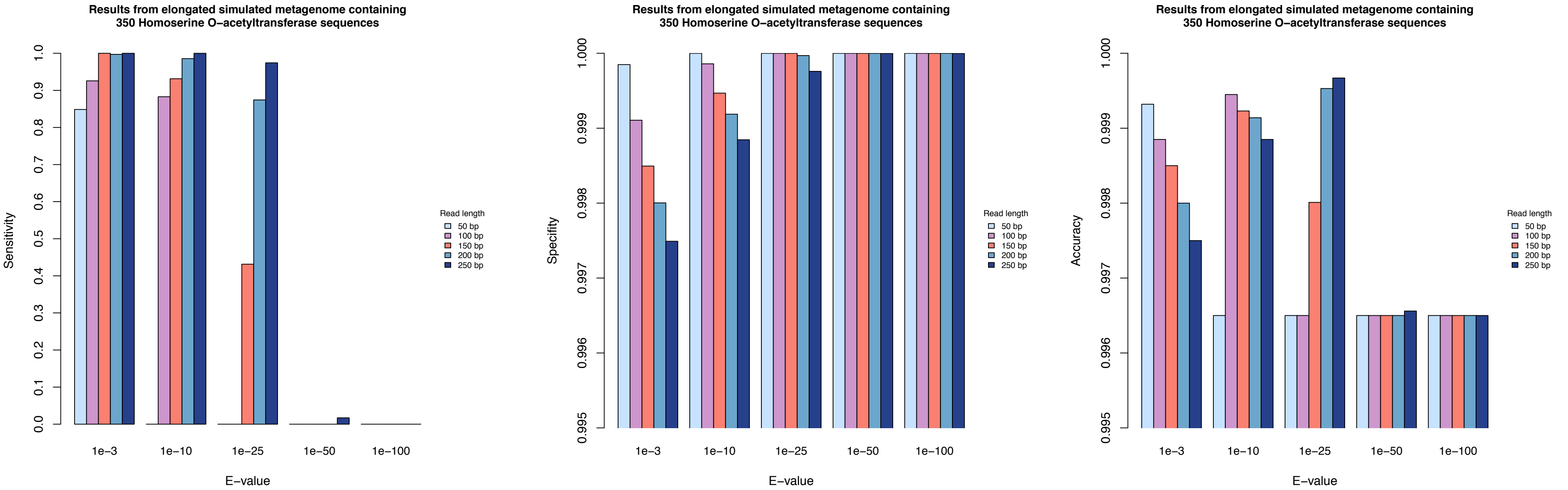
Results from elongated simulated metagenome containing 350 Homoserine O−acetyltransferase sequences
Sensitivity
E−value
1e−3
1e−10
1e−25
1e−50
1e−100
Read length
50 bp
100 bp
150 bp
200 bp
250 bp
Specifity
Accuracy

### Figure S4

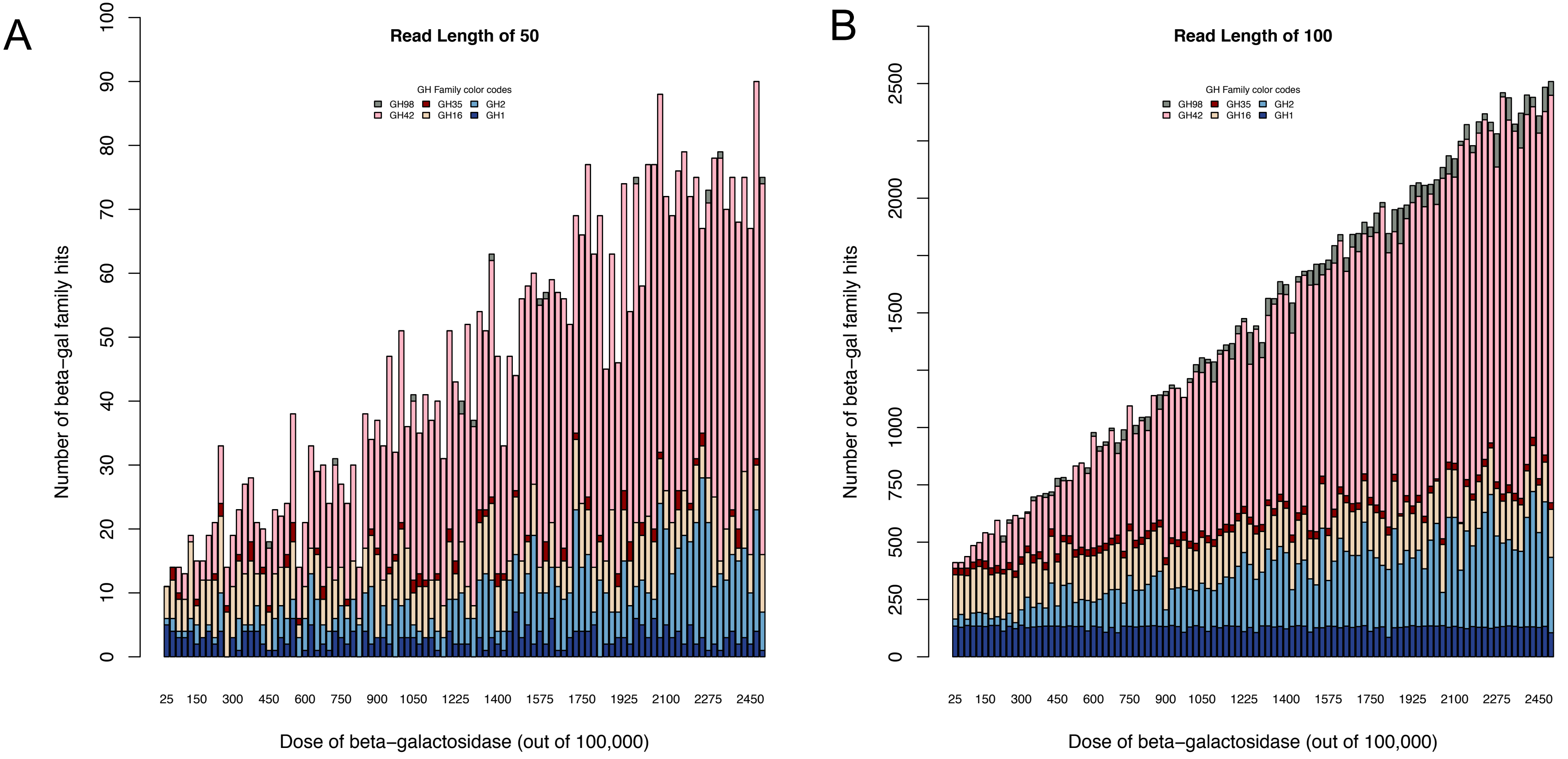


Figure S4: CAZy family distributions across varying beta-gal dosages and read lengths of 50 bp (A), 100 bp (B), 150 bp (C), 200 bp (D), and 250 bp (E). Annotations against 7 CAZy families that contain beta-galactosidase were included: GH1(navy), GH2 (blue), GH16 (tan), GH35 (red), GH42 (pink), GH59 (light green only shown in fig 3D-E), and GH98 (grey). andBased on fig 2A, a read length below 100 bp is inadequate for functional annotation. Generally as read length increases, the overall number annotations to CAZy beta-galactosidase..

Figure S4

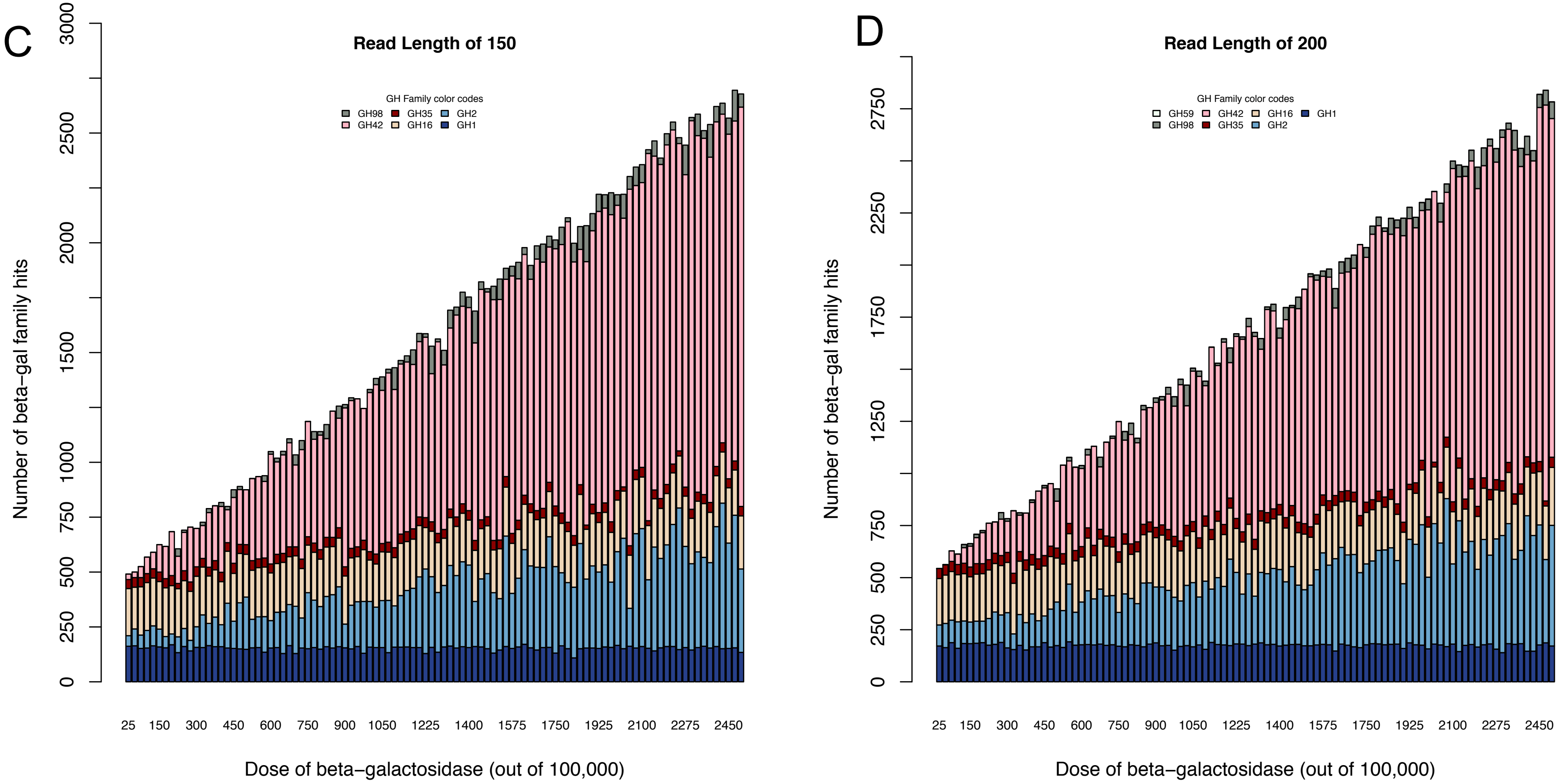


families increases. However, Glycoside Hydrolase Family 1 shows an insignificant increase in annotations with increasing dosages, likely because results include only entire CAZy families, not individual proteins, and matched proteins that are not beta-galactosidase, but in the same family

Figure S4

E

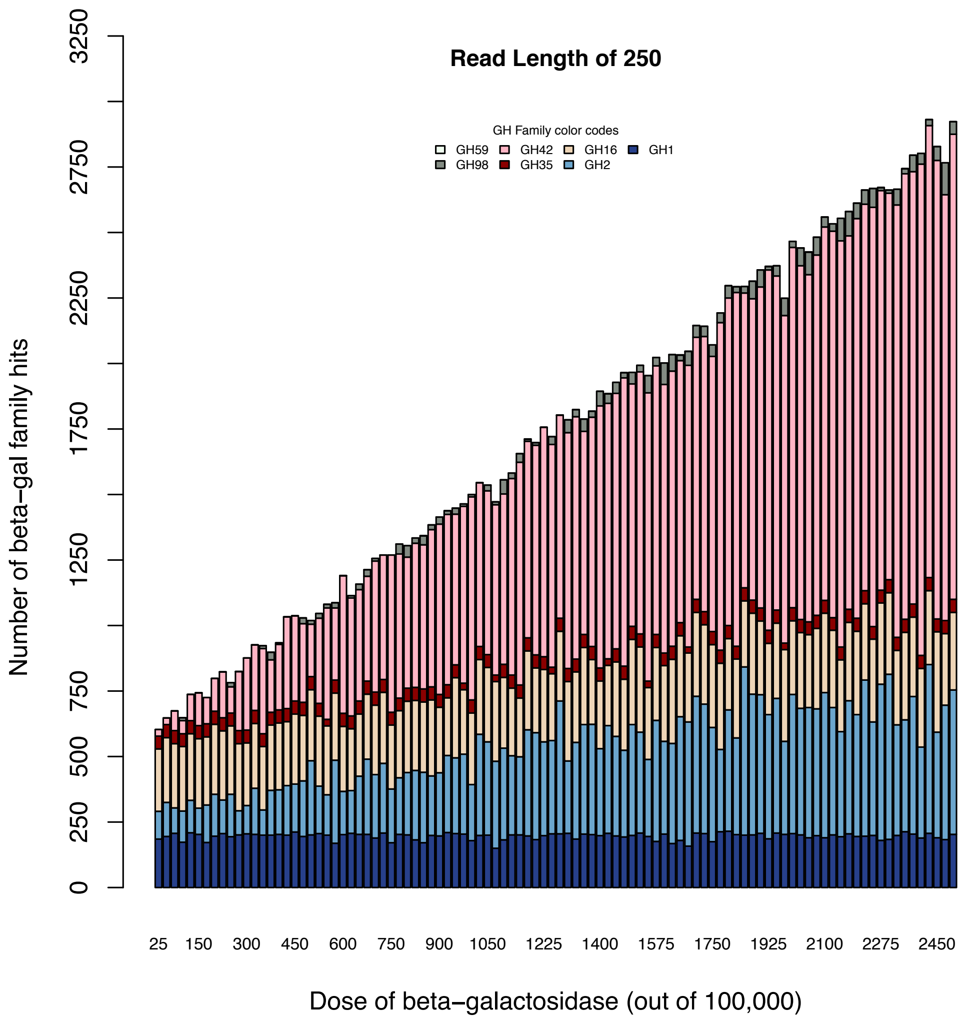

Read Length of 250
GH Family color codes
GH59
GH42
GH16
GH1
GH98
GH35
GH2
Number of beta-gal family hits
0
250
500
750
1250
1750
2250
2750
3250
25
150
300
450
600
750
900
1050
1225
1400
1575
1750
1925
2100
2275
2450
Dose of beta−galactosidase (out of 100,000)

### Figure S5

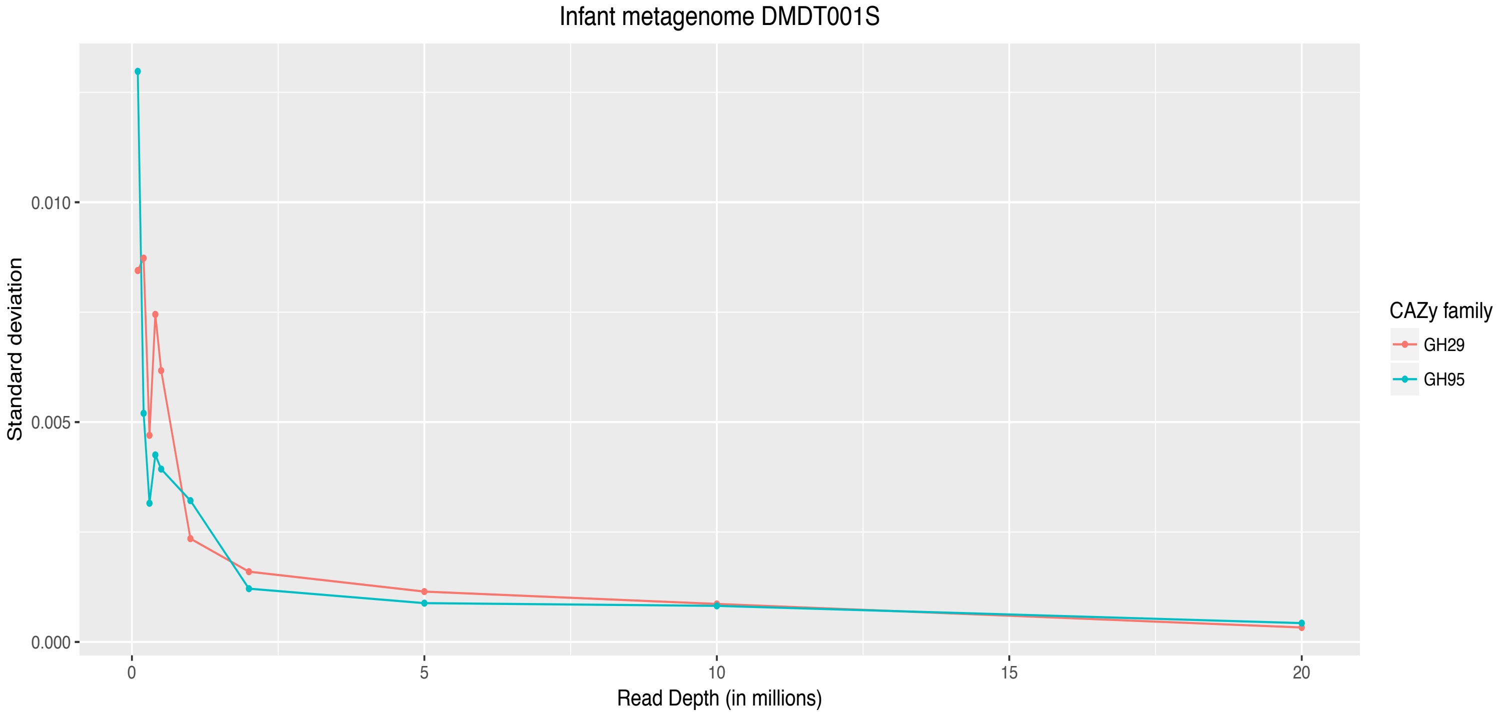


Figure S5: Standard deviation of 10 subsampled metagenomes' annotations against two fucosidase families of the CAZy database for each read depth from an infant metagenome of 30 million merged reads.
